# Testing impacts of goal-oriented outreach with the Girl Scouts: Can a single activity change attitudes towards insects?

**DOI:** 10.1101/2020.12.10.413583

**Authors:** Andrew J. Mongue, Kaila L. Colyott

## Abstract

Most people meet insects with fear and disgust but this reputation is largely unfounded, as few insects pose health risks. In fact, many are beneficial and their absence would adversely affect human life; thus insect conservation is important but unpopular. We have begun addressing these concerns as part of a broader effort to establish an ongoing outreach partnership between graduate students at the University of Kansas and the Girl Scouts of Northeast Kansas/Northwest Missouri. To explore ways to advocate for insect conservation, we held an insect collecting activity at a Girl Scout summer camp and surveyed changes in attitudes towards insects afterwards. This activity positively changed reactions to insect encounters and increased confidence in identifying harmful insects but did not strongly reduce fears or increase curiosity towards insects. Beyond these proximate results, this project highlights the potential of Girl Scout troops as targets for informal science education that can benefit both academics and the broader community.

---

Insects are among the most abundant and diverse groups of organisms, accounting for over half of modern animal life on the planet (Foottit & Adler, 2009). Because of this, humans have encountered insects perhaps more than any other animal (Robinson, 1996), as shown by ancient insect remains among prehistoric settlements (Overgaard Nielsen, Mahler, & Rasmussen, 2000; Panagiotakopulu, 2003). And while we may think that modern ways of life have separated us from natural ecosystems, many insects are very successful in urban environments. Unlike larger animals that need substantial tracts of undisturbed habitats, insects can thrive in small, fragmented urban environments (*e.g.* parks and lawns). And unlike other urban species that are associated with low-income areas (*e.g.* mice, Cohn, Arbes, Yin, Jaramillo, & Zeldin, 2004), insect diversity actually increases in affluent areas (Leong, Bertone, Bayless, Dunn, & Trautwein, 2016), making them a ubiquitous sight in and around homes in all communities.

In spite of, or more likely *because of*, this familiarity, insects are profoundly unpopular with the general public. One study found less than 10% of people enjoy insect encounters in nature and even fewer (<1%) enjoy encountering insects in their home (Byrne, Carpenter, Thoms, & Cotty, 1984). Another survey found that over 85% of people dislike or are afraid of arthropods (the animal phylum including insects, spiders, and crustaceans) found in the house (Hahn & Ascerno, 1991). Some of this fear and disgust may be justified towards disease-vector species, including mosquitoes (Beerntsen, James, & Christensen, 2000; Ledesma & Harrington, 2011) and kissing bugs (Prata, 2001). And researchers have proposed that the disgust that is so often generalized to all insects is an evolutionary behavioral adaptation to avoiding the parasitic or disease-spreading species (Curtis, Aunger, & Rabie, 2004).

Although this instinct may have served our species well historically, it is also demonstrably an overreaction to most commonly encountered insects, the vast majority of which pose no threat to humans. Of those that feed on humans, disease-vectoring is less common than one might imagine: only a small minority (~10%) of mosquito species are known disease transmitters (Rueda, 2008). Moreover, many whole orders of insects such as dragonflies (Odonata) have never been implicated in disease transmission or parasitism of humans. More quantitatively, a survey of biodiversity of arthropods in North Carolina households showed that the majority of species encountered in the American home are benign (Bertone et al., 2016). In other words, much of the generalized dislike of insects is unfounded and ignores many of the beneficial services insects provide.

## A Case for Insects

Non-pest insects play vital roles in ecosystem health and stability, most commonly by breaking down organic matter and facilitating nutrient cycling (Samways, 1994). In forests, for instance, presence of insect herbivores significantly increases available nutrients like nitrogen and phosphorus in the soil (Chapman, Hart, Cobb, Whitham, & Koch, 2003) and up to 20% of wood degradation can be attributed to insects like termites and wood-boring beetles (Ulyshen, 2016). Even in human-made ecosystems, insects fill human-benefiting niches in waste decomposition, like the removal of dung from livestock pastures (Jones & Snyder, 2018), which helps promote grass growth and reduce habitat for insects that parasitize livestock (Fincher, 1981; Gillard, 1967). Even more importantly for the limiting of disease spread, many insects assist in the decomposition of carcasses (Matuszewski, Bajerlein, Konwerski, & Szpila, 2008), a fact that also provides clues to forensic analysts in criminal cases (Buckland & Smith, 1988; Byrd, 2002).

Finally, and most popularly known, insects also pollinate many plants, including agricultural crops. The majority of crops are at least partially insect-dependent for pollination and fruit production, and crops like almonds, hay, and blueberries are completely dependent on insect pollinators (Morse & Calderone, 2000). The economic value of bee pollination alone in the United States provides services worth upwards of $5 billion (Southwick & Southwick, 1992). Add to this the other benefits, including those described above, and the total value of all insect services to society is estimated at $57 billion in the United States alone (Losey & Vaughn, 2006). Thus, the overwhelming fear and dislike of insects and their relatives is not only unfounded but also problematic from an economic point of view.

## Generating Public Support

For all of the above reasons, there is a great need to promote acceptance and conservation of insects, but little has been done to advocate for this group of animals. Most conservation efforts focus on charismatic species, typically large mammals (e.g. whales, Scott & Parsons, 2005). Only a few well-known insects, most prominently the monarch butterfly, have received comparable attention (Diffendorfer et al., 2014; Missrie & Nelson, 2005; Oberhauser & Solensky, 2004). Given the scale of insect diversity, rather than attempting to generate case-by-case popularity, a greater ecosystem-level and, indeed, human benefit could be obtained with conservation of the broader group of arthropods, focusing on their positive contribution as members of an ecological community (Hughes, Daily, & Ehrlich, 2000; Panzer & Schwartz, 1998; Samways, 2007). Before tackling more comprehensive conservation efforts however, public attitude towards insects must be improved to ensure the success of those efforts. In this study, we sought to quantify how effective single-intervention teaching is in changing attitudes and reactions towards insects. We focused on a demographic with perhaps the worst perceptions of insects: grade-school-aged girls.

## Outreach Partnership with the Girl Scouts

Our efforts to change insect popularity grew from a broad partnership with the Girl Scouts of Northeast Kansas/Northwest Missouri. After initial successful volunteer events with local troops, we were approached by Girl Scout program managers to expand involvement and increase outreach teaching opportunities for graduate students at the University of Kansas. With a formal community partnership, graduate students designed five single-activity modules based on both their research interests and relevance to teaching objectives for Girl Scout badges. These activities were hosted on the Girl Scouts’ community partner webpage and troop leaders could then contact these graduate students to schedule an activity for their troop. This arrangement benefitted both parties, as troop leaders could select the most relevant activity for the needs or interests of their girls and graduate students offered activities most directly relevant to their own interests and expertise. Over the course of two years, these programs have reached roughly 500 girls and resulted in our programs receiving a 2018 Community Collaboration Award.

For a specific example, we, the authors, began by offering an activity to help Brownie Scouts meet requirements to earn their Bugs Badge. The badge has multiple components ranging from insect themed arts and crafts to exploration of insect habitats. We focused on the latter, showing girls where and how to collect local insects. This activity was one of the more popular, being requested by 166 girls in total. Owing to the qualitative change in attitudes we noticed in girls who participated in these activities in the first year of our partnership, we designed a simple survey to test whether a single activity interacting with arthropods could reduce fear and increase appreciation of local insect species.

We obtained Institutional Review Board approval for study design and consent language from the University of Kansas (IRB ID: 00141007) and carried out the survey at a Girl Scout summer camp in the summer of 2017. We informed the parent or guardian of each participating child upon their arrival at camp that their child was in a camp group connected to a research study. We gave the parent or guardian a verbal summary of the project and a paper copy of the survey to review before asking them to sign a consent form allowing their child to participate in the study. Parents had the option of opting out of the study by not signing the consent form, without affecting their child’s ability to participate in camp activities, including insect collection. Children without parental or guardian consent were not given a survey to complete, and no identifying information was collected for any child during the survey process.

We administered the survey to groups of Junior and Cadette rank girls for two months in the summer of 2017 at Camp Daisy Hindman, in rural Dover, Kansas (n = 88 total respondants). To minimize identifiable information collected, we did not record ages of participants, but these ranks correspond to 4^th^ to 8^th^ grade students. Throughout the summer camp season (June – July), we visited the camp each week and collected data from two groups of girl. Each week, one group worked with us on an insect collecting activity before taking a survey of attitudes and reactions towards insects. The second (control) group had no interaction with us prior to the survey. Collecting activities varied by week (blacklight trapping, stream collecting, or open field sweep netting) depending on the camp program and weather, but in each activity girls collected insects and transfered them from a net to a mesh cage by hand. Throughout the activity, we encouraged girls to share their discoveries and help each other with collecting. With the girls’ consent, we saved representative specimens of collected species to be pinned and spread by us as part of a display kept at the camp.

For each activity, we used a teaching collection of pinned insects to facilitate a short discussion (~10 minutes) that included an overview of stinging insects and an open-ended question and answer session. Immediately after the hands-on collecting session, we spent a short time (~5 minutes), asking girls to share their favorite catches. The girls for whom we had obtained prior parental consent were then given a survey to fill out. For the control group, girls were given surveys immediately after completing their regularly scheduled camp activities (*e.g.* tie-dying or horseback riding) with no collecting activity or discussion of insects. Camp groups that were chosen as control groups were selected to keep the number of participants and ages roughly equal between the treatment and control.

## Survey Content and Analysis

The anonymous surveys consisted of 15 questions, with 3 background questions and 12 retrospective before/after questions that asked participants to answer how they felt both before and after their time at the summer camp (full survey shown in Table 1). The use of a retrospective pretest-posttest design (*i.e*. administering both the pre- and post-test questions after the intervention) provides a more accurate assessment of change than a conventional pretest-posttest design (*i.e*. administering pre-test before and post-test after), because it allows the respondent to use a consistent scale when answering questions about both the present and past (Nakonezny & Rodgers, 2005).

**Table 1.** The survey questions presented to girls and analyzed here. Participants were asked to circle an answer to each question either after the insect activity (treatment) or immediately upon gathering (controls).

|  |  |  |  |  |
| --- | --- | --- | --- | --- |
| **1. Did you spend time working with the bug people at camp?** | **YES** | **NO** |  |  |
| **2. How often do you encounter bugs at home?** | Always | Often | Sometimes | Never |
| **3. How often do you encounter spiders at home?** | Always | Often | Sometimes | Never |
| **4. How afraid were you when encountering bugs at home BEFORE coming camp?** | Very afraid | Somewhat | Not very | Not afraid |
| **5. How afraid were you when encountering spiders at home BEFORE coming camp?** | Very afraid | Somewhat | Not very | Not afraid |
| **6. BEFORE coming to camp, when you encountered a bug at home, what would you do?** | Kill it | Run away | Ignore it | Move it outside |
| **7. BEFORE coming to camp, when you encountered a spider at home, what would you do?** | Kill it | Run away | Ignore it | Move it outside |
| **8. How likely were you to pick up a bug and be curious about it at home, BEFORE coming to camp?** | Always | Often | Sometimes | Never |
| **9. How good do you think you were at determining if a bug was dangerous or not, BEFORE coming to camp?** | Great | Good | OK | Not good |
| **10. How afraid are you of encountering bugs at home AFTER coming camp?** | Very afraid | Somewhat | Not very | Not afraid |
| **11. How afraid are you of encountering spiders at home AFTER coming camp?** | Very afraid | Somewhat | Not very | Not afraid |
| **12. AFTER coming to camp, when you encountered a bug at home, what will you do?** | Kill it | Run away | Ignore it | Move it outside |
| **13. AFTER coming to camp, when you encountered a spider at home, what will you do?** | Kill it | Run away | Ignore it | Move it outside |
| **14. How often do you think you will pick up a bug and be curious about it, AFTER coming to camp?** | Always | Often | Sometimes | Never |
| **15. How good do you think you are at determining if a bug is dangerous or not, AFTER coming to camp?** | Great | Good | OK | Not good |

Most questions were based on a Likert-like scale of responses (*e.g.* always / often / sometimes / never) but the reactions to the encounter questions were subjectively ranked from least desirable to most: killing the insect or spider (fearful and destructive), running away (fearful and passive), ignoring it (neutral/non-destructive), moving it outside (active and unafraid). Background questions were implemented as a check to ensure no systematic differences existed in everyday exposure to arthropods between our treatment and control groups. The twelve retrospective before/after questions were also designed in pairs for control: one set asked about attitudes towards insects (“bugs” in the survey) and the other asked about spiders, which were not a part of the hands-on teaching or open-ended discussion. This pairing created an additional check that time spent at camp was not changing attitudes about invertebrates in general by virtue of bringing campers closer to nature than they would be at home.

Prior to downstream analyses, we curated the data for irregularities. A small minority of girls skipped questions, chose multiple answers to a single question, or answered in a manner seemingly contrary to the design of the experiment (*i.e.* individuals from the control group indicating that they worked with us, despite no interaction at camp prior to the survey). This last class of problems was rare but potentially confounding, as we had been doing community outreach workshops for the two preceding years in the area, so some girls in our control groups may have had previous experience with our teaching outside the scope of this project. To be conservative in analyses, we discarded both of the surveys that had the uncertain treatment status described above; this curation brought our sample size down from 88 to 86 (45 treatment, 41 control). For the remaining irregularities, answers were excluded on a per-case basis (*e.g*. a girl who skipped or gave multiple answers to question 3 would still have her answers to 4-15 included in analyses), resulting in slight differences in sample sizes between questions. We coded each potential response to a question as a number from 0 to 3 for analysis. While these data are not continuous and not necessarily normally distributed, parametric tests should be robust to these assumptions given our sample sizes surveyed. Thus we assessed simple differences in the treatment and control groups with t-tests for the background questions.

To parse the more complicated effect of our lessons while controlling for time at camp, we opted to analyze results in a permutation framework that made no assumptions about underlying data distributions. First we calculated the difference in means before and after time at camp in the treatment and control groups separately. Then we calculated the difference of these differences to get a measure of how dissimilar the treatment and control groups were while controlling general experiences at camp. To assess significance of these differences, we then carried out permuations randomly assigning girls to treatment or control groups of sizes equal to the true groups. As before we calculated the difference of differences between our pseudo-treatment and pseudo-control groups. By repeating this for one thousand permutations, we generated an expectation of differences between groups which could occur by chance. We then compared our true value to this distribution; the p-value was taken as the proportion of times the the true difference was more extreme than the randomly generated differences. As such, there are no test-statistics *per se* to report for these analyses, only p-values. All analyses and data visualizations were carried out using custom scripts written in R version 3.4.1(R Core Team, 2017).

## Findings

The control and treatment groups did not differ in exposure to insects at home (Question 2; t_78.1_ = 0.19, p = 0.85) but, oddly, they reported a difference in spider enounters with the control group encountering fewer spiders (Question 3; t_80.5_ = 2.41, p = 0.02). This starting difference is less relevant for our focus on insects, and moreover appears to have no biasing effect, as groups did not differ from each other in their reaction to (p = 0.555, Questions 7 & 13) or fear of (p = 0.293, Questions 5 & 11) spiders while controlling for time at camp.

With regard to insects, two of our metrics showed significant changes in our treatment group after the activity. Girls became more confident in being able to identify dangerous insects (p = 0.018, Questions 9 & 15) and became less likely to kill or run away from an insect encountered at home (p = 0.041, Questions 6 & 12). Results with sample sizes can be seen in Figure 1A and B respectively. Our two other metrics, curiousity (Questions 8 & 14) and fear of insects (Questions 4 & 10) did not significantly change after our lesson, but did trend in the direction of more curiosity (p = 0.099) and less fear (p = 0.180). In the latter case, both treatment and control groups reported marginal decreases in fear after their time at camp. These results, including sample sizes, are summarized in Figure 2A and B.

**Figure 1.**
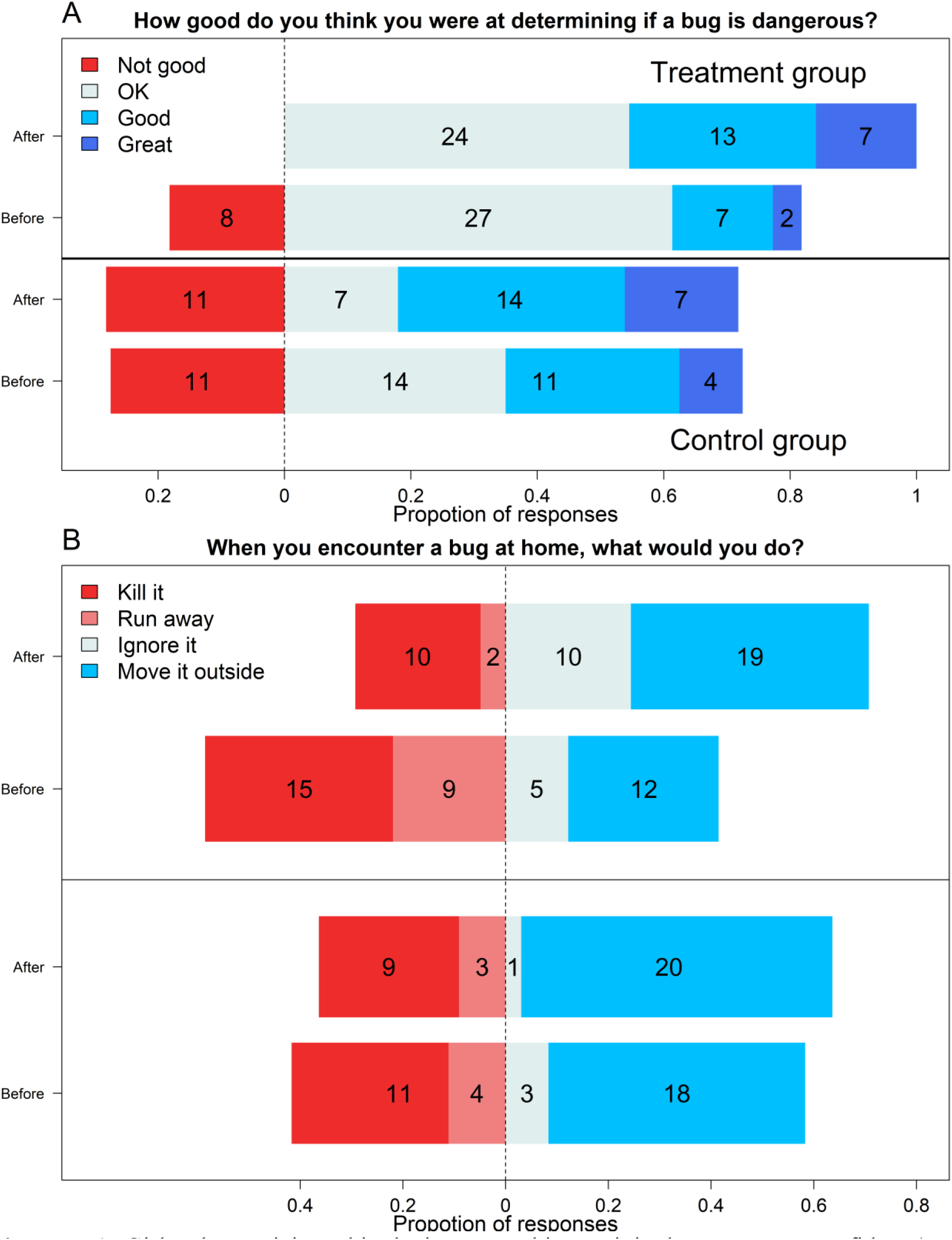
**A**. Girls who participated in the insect catching activity became more confident (p = 0.018) in identifying dangerous insects (top panel) than their control counterparts (bottom panel). **B.** Likewise, girls from the treatment group became more likely to move an insect outside and less likely to kill it or run away (p = 0.041) after the lesson. Each response category is color-coded per the legend and numbers in each category represent counts of respondents.

**Figure 2.**
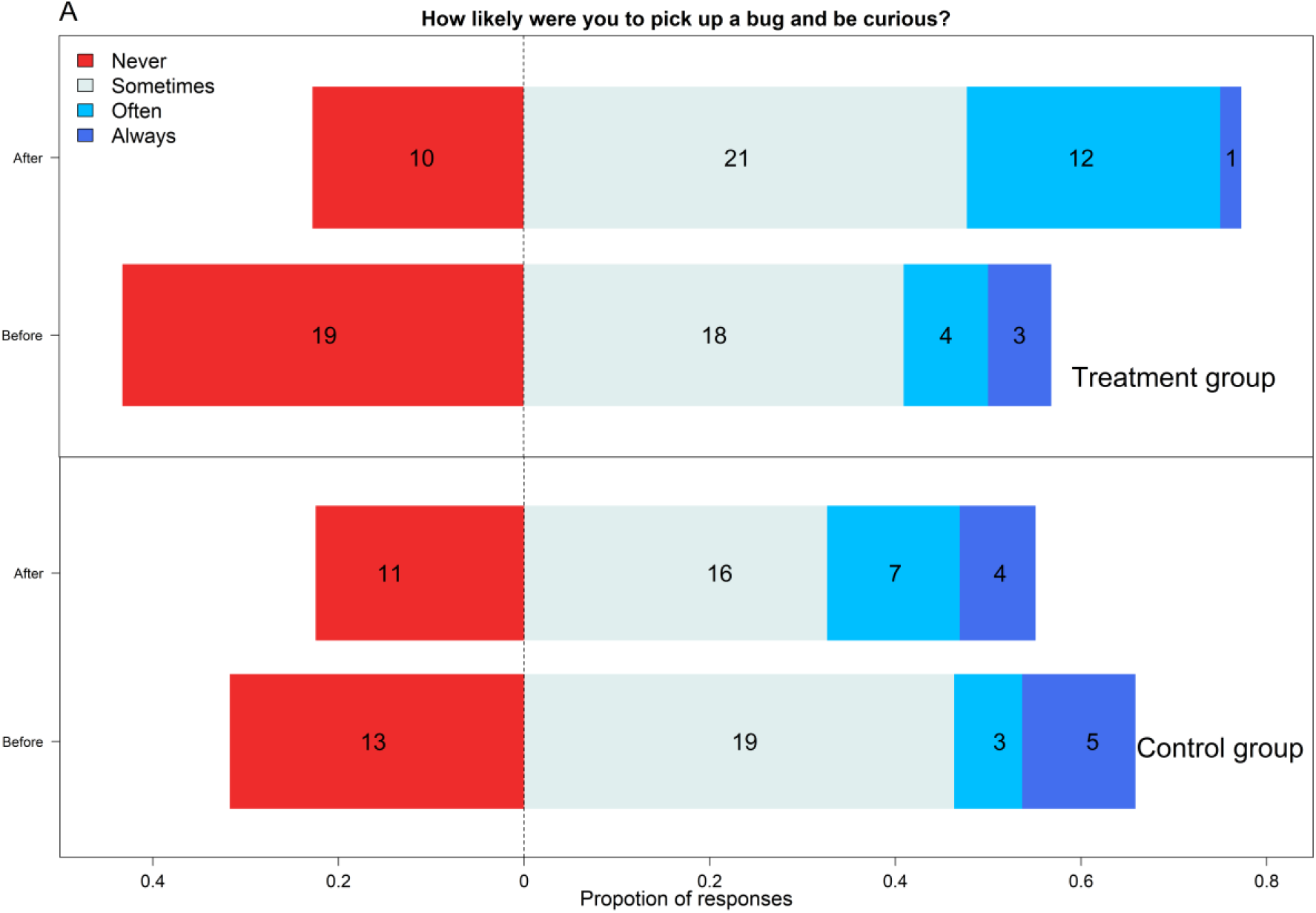
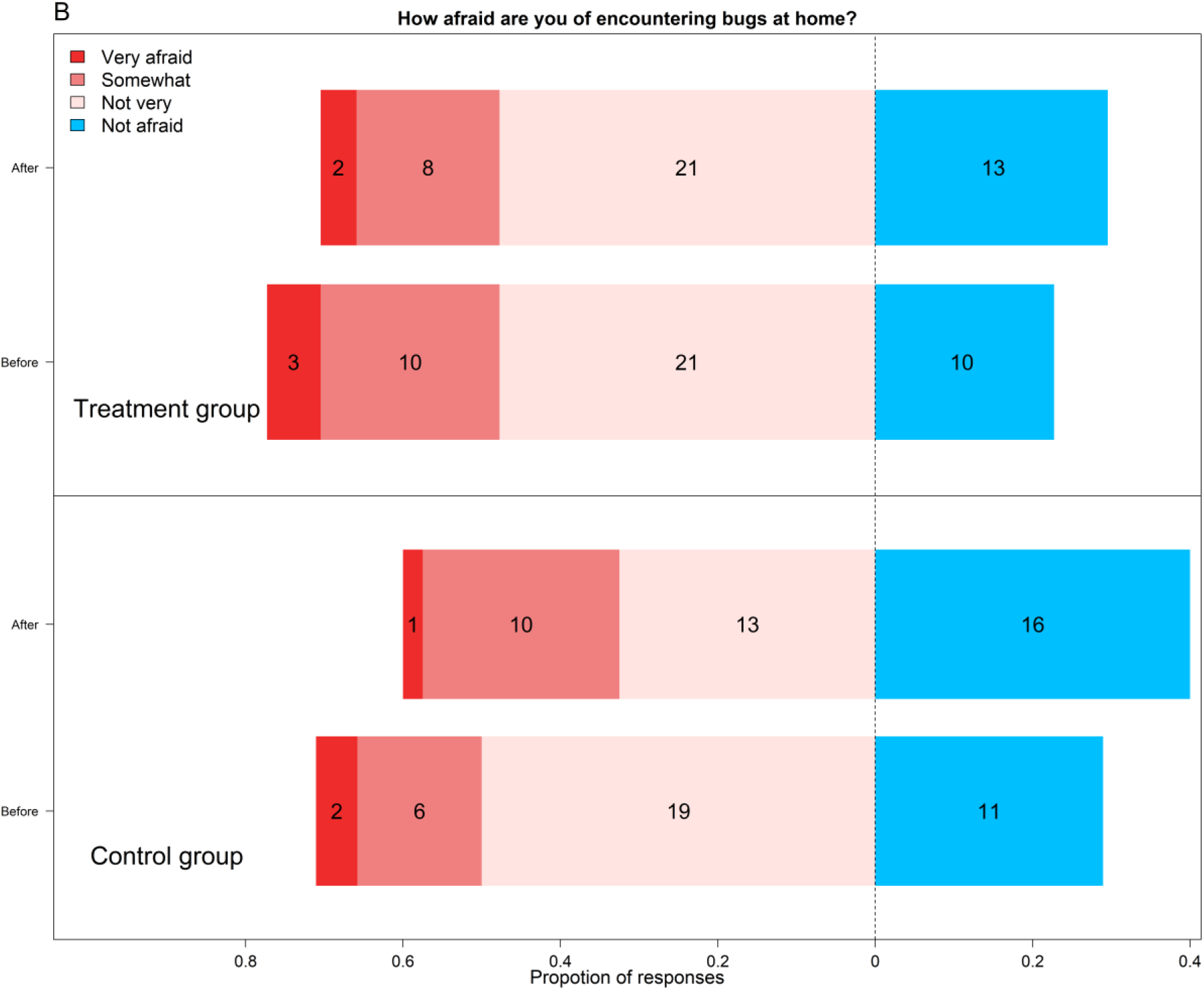
**A.** Changes in girls’ curiosity and willingness to pick up insects for both treatment and control groups. The difference in the two groups after treatment was not significant (p = 0.099), but trended towards an increase in curiosity as can be seen in the top panel. **B.** Our activity did not decrease respondents’ fears of insects (p = 0.180) in any meaningful way. As with the other figure, each response category is color-coded per the legend and numbers in each category represent counts of respondents.

## Discussion

The public in general dislikes insects more than most animals (Byrne et al., 1984) and women more than men find them disgusting (Curtis et al., 2004). In a targeted effort to change these attitudes, we designed a set of activities to carry out with the girls of Girl Scout Camp Daisy Hindman. Afterwards, we surveyed girls to assess how effective activities involving live insects can be in changing perceptions. In brief, we found that we increased confidence in differentiating dangerous and harmless insects and positively changed self-reported reactions to an insect encounter. Girls became less likely to kill insects encountered in the home and more likely to move them outside or ignore them. Both of these results are encouraging for the goal of increasing acceptance of insects. Obviously decreasing the instinct to immediately kill an insect found in the home can only help conserve insects, and learning to confidently differientiate dangerous and harmless insects should lead to fewer perceived threats from encounters with harmless insects. Whether or not this effect persists in the longterm would be an obvious target for future study.

On the other hand, we failed to strongly increase curiosity or decrease fear of insects in the span of this activity. However, both of these metrics showed small changes in the desired direction after our lesson, so it is possible that this hands-on approach could be effective but would require more engagement time to generate strong changes. As with the positive results, it would also be fruitful to examine the fear and curiosity components after repeated interactions.

Another possibility is that we failed to identify fears in a precise enough way to notice a change. Girls who participated reported becoming more confident in identifying dangerous insects but did not report a decrease in fear of “bugs” as a blanket category. Considering these two outcomes together, it would be interesting to ask about fears of specific groups of insects to see if fears become less generalized and more concentrated on groups that can cause harm like ants and wasps. We avoided such detailed questions in this initial survey out of a desire to keep the survey portion short and easy to complete but it would be appropriate for a more targeted follow-up study.

More generally, our activity sparked engagement in spite of using an unpopular group of animals suggesting a great potential to stimulate excitement with this age group. The surveyed scouts were late elementary school to middle school aged, the time when girls become less likely than boys to pursue interests in sciences (Clark Blickenstaff, 2005), so similarly hands-on approaches with Girl Scouts make an obvious target for promoting women in STEM (science, technology, engineering, and math).

## Lessons Learned: Graduate-Student-Girl-Scout Partnerships as a Mutually Beneficial Relationship

We found that our regional Girl Scout organization offers a receptive audience for informal STEM education and we suggest that they make an excellent venue for outreach across the sciences. By advertising our STEM expertise and taking requests for outreach teaching, we were able to match our science skillset with a demand in the community. This two-way interaction sparked our initial interest in formally assessing the effectiveness of our outeach activities and we submit that this approach can serve as a useful model for goal-oriented outreach among academic researchers.

Although such outreach may be more common among other educational groups, for many research-focused scientists, outreach remains an unorganized endeavor. Developing broad community impacts is an important component of many acadedmic positions, but it often receives less attention than research or formal (*i.e.* classroom-based) teaching. We submit that outreach can and should be approached in the same manner as the rest of the scientific process: with concrete objectives and empirical validation to assess how successfully these objectives are met. Under this paradigm, outreach events are more beneficial to both the researchers and the public. Researchers can have meaningful interactions and encourage interest in science as we saw with girls’ confidence in insect identification and decreased instinct to kill insects in this study.

## Next Steps

The established structure and persistent groups of Girl Scout troops make excellent targets for repeated scientific engagement across multiple years even. Ancedotally, we have seen some girls at multiple outreach events, but, due to our limited time in graduate school, we are not able to formally track the longer term impacts of our activities on either interest in science or attitudes towards insects. While we, the authors, have since graduated, we are happy to report that the Girl Scout partnership still exists with current graduate students at the University of Kansas and continues to offer a platform for informal STEM teaching. In its current incarnation, the partnership consists of the continued independent modules as well as an annual STEM activity day at one of the camps (Camp Tongawood); current graduate students have plans to use this venue for outreach outcome surveying.

## Acknowledgments

We would like to thank all those who made this work possible, including the girls and parents who participated in this survey, the executives of the Girl Scouts of Northeast Kansas/Northwest Missouri, and all of the counsellors and coordinators at Camp Daisy Hindman, especially the camp director, Marley Parsons, who was always the first to help us coordinate our efforts. Thanks to Andrea Lucky for encouraging us to publish, Jennifer Gleason for sponsoring our IRB project, and the other members of the Ecology and Evolutionary Biology Outreach Committee for support throughout.

## References

Beerntsen, B. T., James, A. A., & Christensen, B. M. (2000). Genetics of Mosquito Vector Competence. Microbiology and Molecular Biology Reviews, 64(1), 115–137. https://doi.org/10.1128/MMBR.64.1.115-137.2000

Bertone, M. A., Leong, M., Bayless, K. M., Malow, T. L. F., Dunn, R. R., & Trautwein, M. D. (2016). Arthropods of the great indoors: Characterizing diversity inside urban and suburban homes. PeerJ, 4, e1582. https://doi.org/10.7717/peerj.1582

Buckland, P. C., & Smith, K. G. V. (1988). A Manual of Forensic Entomology. American Journal of Archaeology, 92(2), 287. https://doi.org/10.2307/505635

Byrd, J. H. (2002). Forensic entomology: The utility of arthropods in legal investigations. CRC press.

Byrne, D. N., Carpenter, E. H., Thoms, E. M., & Cotty, S. T. (1984). Public attitudes toward urban arthropods. Bulletin of the ESA, 30(2), 40–44.

Chapman, S. K., Hart, S. C., Cobb, N. S., Whitham, T. G., & Koch, G. W. (2003). Insect herbivory increases litter quality and decomposition: An extension of the acceleration hypothesis. Ecology, 84(11), 2867–2876.

Clark Blickenstaff*, J. (2005). Women and science careers: Leaky pipeline or gender filter? Gender and Education, 17(4), 369–386.

Cohn, R. D., Arbes, S. J., Yin, M., Jaramillo, R., & Zeldin, D. C. (2004). National prevalence and exposure risk for mouse allergen in US households. Journal of Allergy and Clinical Immunology, Vol. 113, pp. 1167–1171. https://doi.org/10.1016/j.jaci.2003.12.592

Curtis, V., Aunger, R., & Rabie, T. (2004). Evidence that disgust evolved to protect from risk of disease. Proceedings of the Royal Society of London. Series B: Biological Sciences, 271(suppl_4), S131–S133.

Diffendorfer, J. E., Loomis, J. B., Ries, L., Oberhauser, K., Lopez-Hoffman, L., Semmens, D.,… Thogmartin, W. E. (2014). National valuation of monarch butterflies indicates an untapped potential for incentive-based conservation. Conservation Letters, 7(3), 253–262. https://doi.org/10.1111/conl.12065

Fincher, G. T. (1981). The potential value of dung beetles in pasture ecosystems. Journal of the Georgia Entomological Society, 16, 316–333. https://doi.org/10.1094/PDIS-05-17-0721-RE

Foottit, R. G., & Adler, P. H. (2009). Insect Biodiversity: Science and Society. In Insect Biodiversity: Science and Society. https://doi.org/10.1002/9781444308211

Gillard, P. (1967). Coprophagous beetles in pasture ecosystems. Journal of the Australian Institute of Agricultural Science, 33(1), 30–34 pp.

Hahn, J. D., & Ascerno, M. E. (1991). Public Attitudes Toward Urban Arthropods. American Entomologist, (fall), 179–185. https://doi.org/10.1093/ae/37.3.179

Harcourt, A. H. (1986). Gorilla conservation: Anatomy of a campaign. In Primates (pp. 31–46). Springer.

Hughes, J. B., Daily, G. C., & Ehrlich, P. R. (2000). Conservation of insect diversity: A habitat approach. Conservation Biology, 14(6), 1788–1797. https://doi.org/10.1046/j.1523-1739.2000.99187.x

Jones, M. S., & Snyder, W. E. (2018). Beneficial Insects in Agriculture: Enhancement of Biodiversity and Ecosystem Services. Insect Biodiversity: Science and Society, 2, 105–122.

Ledesma, N., & Harrington, L. (2011). Mosquito vectors of dog heartworm in the United States: Vector status and factors influencing transmission efficiency. Topics in Companion Animal Medicine, 26(4), 178–185. https://doi.org/10.1053/j.tcam.2011.09.005

Leong, M., Bertone, M. A., Bayless, K. M., Dunn, R. R., & Trautwein, M. D. (2016). Exoskeletons and economics: Indoor arthropod diversity increases in affluent neighbourhoods. Biology Letters, 12(8), 23. https://doi.org/10.1098/rsbl.2016.0322

Losey, J. E., & Vaughn, M. (2006). The economic value of ecological services provided by insects. BioScience, 56(4), 311–323. https://doi.org/10.1641/0006-3568(2006)56[311:TEVOES]2.0.CO;2

Matuszewski, S., Bajerlein, D., Konwerski, S., & Szpila, K. (2008). An initial study of insect succession and carrion decomposition in various forest habitats of Central Europe. Forensic Science International, 180(2-3), 61–69. https://doi.org/10.1016/j.forsciint.2008.06.015

Missrie, M., & Nelson, K. (2005). Direct payments for conservation: Lessons from the Monarch Butterfly Conservation Fund. Economics, 3(88), 339–353.

Morse, R. A., & Calderone, N. W. (2000). The value of honey bees as pollinators of US crops in 2000. Bee Culture, 128(March 2000), 1–15.

Nakonezny, P. A., & Rodgers, J. L. (2005). An empirical evaluation of the retrospective pretest: Are there advantages to looking back? Journal of Modern Applied Statistical Methods, 4(1), 22.

Oberhauser, K. S., & Solensky, M. J. (2004). The Monarch butterfly: Biology & conservation. Cornell university press.

O’Connell-Rodwell, C. E., Rodwell, T., Rice, M., & Hart, L. A. (2000). Living with the modern conservation paradigm: Can agricultural communities co-exist with elephants? A five-year case study in East Caprivi, Namibia. Biological Conservation, 93(3), 381–391.

Overgaard Nielsen, B., Mahler, V., & Rasmussen, P. (2000). An arthropod assemblage and the ecological conditions in a byre at the Neolithic settlement of Weier, Switzerland. Journal of Archaeological Science, 27(3), 209–218. https://doi.org/10.1006/jasc.1999.0448

Panagiotakopulu, E. (2003). Insect remains from the collections in the Egyptians Museum of Turin. Archaeometry, 45(2), 355–362. https://doi.org/10.1111/1475-4754.00113

Panzer, R., & Schwartz, M. W. (1998). Effectiveness of a Vegetation-Based Approach to Insect Conservation Effectiveness to Insect of a Approach Conservation. Conservation Biology, 12(3), 693–702. https://doi.org/10.1046/j.1523-1739.1998.97051.x

Prata, A. (2001). Clinical and epidemiological aspects of Chagas disease. The Lancet. Infectious Diseases, 1(2), 92–100. https://doi.org/10.1016/S1473-3099(01)00065-2

R Core Team. (2017). R Core Team (2017). R: A language and environment for statistical computing. R Foundation for Statistical Computing, Vienna, Austria. URL http://www.r-project.org/., R Foundation for Statistical Computing.

Robinson, W. H. (1996). Urban entomology: Insect and mite pests in the human environment. Chapman & Hall.

Rueda, L. M. (2008). Global diversity of mosquitoes (Insecta: Diptera: Culicidae) in freshwater. Hydrobiologia, 595(1), 477–487.

Samways, M. J. (1994). Insect Conservation Biology (Conservation Biology, No 2). Springer Science & Business Media.

Samways, M. J. (2007). Insect Conservation: A Synthetic Management Approach. Annual Review of Entomology, 52(1), 465–487. https://doi.org/10.1146/annurev.ento.52.110405.091317

Scott, N. J., & Parsons, E. C. M. (2005). A survey of public opinion in south-west Scotland on cetacean conservation issues. Aquatic Conservation: Marine and Freshwater Ecosystems, Vol. 15, pp. 299–312. https://doi.org/10.1002/aqc.662

Southwick, E. E., & Southwick, L. (1992). Estimating the Economic Value of Honey Bees (Hymenoptera: Apidae) as Agricultural Pollinators in the United States. Journal of Economic Entomology, 85(3), 621–633. https://doi.org/10.1093/jee/85.3.621

Ulyshen, M. D. (2016). Wood decomposition as influenced by invertebrates. Biological Reviews, 91(1), 70–85. https://doi.org/10.1111/brv.12158

Zuo, J., Cornelissen, J. H. C., Hefting, M. M., Sass-Klaassen, U., van Logtestijn, R. S. P., van Hal, J.,… Berg, M. P. (2016). The (w)hole story: Facilitation of dead wood fauna by bark beetles? Soil Biology and Biochemistry, 95, 70–77. https://doi.org/10.1016/j.soilbio.2015.12.015

